# SERS uncovers the link between conformation of cytochrome *c* heme and mitochondrial membrane potential

**DOI:** 10.1101/2021.01.03.425119

**Authors:** Nadezda A. Brazhe, Evelina I. Nikelshparg, Adil A. Baizhumanov, Vera G. Grivennikova, Anna A. Semenova, Sergey M. Novikov, Valentyn S. Volkov, Aleksey V Arsenin, Dmitry I. Yakubovsky, Andrey B. Evlyukhin, Zhanna V. Bochkova, Eugene A. Goodilin, Georgy V. Maksimov, Olga Sosnovtseva, Andrey B. Rubin

## Abstract

The balance between the mitochondrial respiratory chain activity and the cell’s needs in ATP ensures optimal cellular function. Cytochrome *c* is an essential component of the electron transport chain (ETC), which regulates ETC activity, oxygen consumption, ATP synthesis and can initiate apoptosis. The impact of conformational changes in cytochrome *c* on its function is not understood for lack of access to these changes in intact mitochondria. We have developed a novel sensor that uses unique properties of label-free surface-enhanced Raman spectroscopy (SERS) to identify conformational changes in heme of cytochrome *c* and to elucidate their role in functioning mitochondria. We verify that molecule bond vibrations assessed by SERS is a reliable indicator of the heme conformation during changes in the inner mitochondrial membrane potential and ETC activity. We have found that cytochrome *c* heme reversibly switches between planar and ruffled conformations in response to the inner mitochondrial membrane potential and H^+^ concentration in the intermembrane space to regulate the efficiency of the mitochondrial respiratory chain, thus, adjusting the mitochondrial respiration to the cell’s consumption of ATP and the overall activity. The ability of the proposed SERS-based sensor to track mitochondrial function opens wide perspectives on cell bioenergetics.

**For Table of Contents Only:** 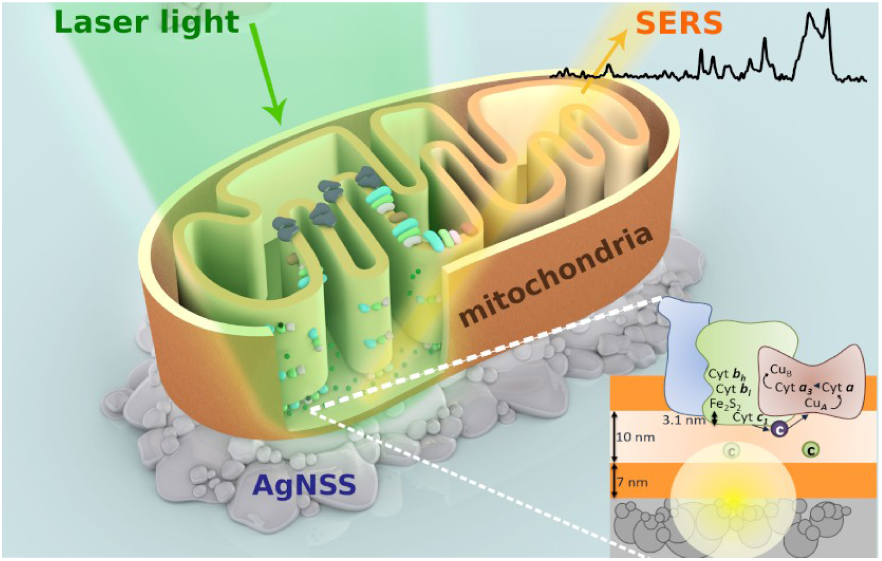

---

Mitochondrial respiratory chain (electron transport chain, ETC) provides oxidative phosphorylation (OXPHOS) and plays a crucial role in the production of reactive oxygen species (ROS), initiation of apoptosis, maintenance of Ca^2+^ concentration in cytoplasm and its storage^1,2,3^. ETC activity determines the efficiency of ATP synthesis. An imbalance between ETC activity and the amount of ATP needed by the cell may lead to ATP deficiency and suppression of the cell function, or to the excessed ROS production in ETC, oxidative stress, and cytochrome *c*-dependent apoptosis^4,5,6,7^. To eliminate these negative effects mitochondria possess various regulatory mechanisms to adjust the activity of ETC complexes to the optimal rate of ATP synthesis^8,9^.

Cytochrome *c* (cytC) is a small globular protein containing heme *c* and represents the only soluble electron carrier in ETC, transferring electrons from *bc1*-complex (complex III) to cytochrome *c* oxidase (complex IV) by means of diffusion in the mitochondrial intermembrane space (IMS)^10,11^. Tissue-specific phosphorylation of cytC affects its activity and the whole OXPHOS^12,13,14,15^. Huttemann et al. suggested that high membrane potentials (ΔΨ) drive cytC phosphorylation in order to prevent inner mitochondrial membrane (IMM) hyperpolarization and ROS production^12^. Growing evidence indicates that cytC activity can be regulated not only by the modification of the cytC protein part but also by heme conformational changes^16^. It is known that cytC heme predominantly exists in two conformations: planar and ruffled^17^. Purified wild type cytC and its various mutant forms exhibit different probabilities for planar and ruffled heme conformations; cytC mutants with the ruffled heme have a decreased rate of electron transfer between cytC – complex III and cytC – complex IV^17,18,19^. Experiments on the solution of purified oxidized cytC under high electric fields also revealed ruffled heme conformation^20^. In mitochondria, the inner mitochondrial membrane potential might affect heme conformation and cytC ability to transfer electrons. This is unknown yet for lack of techniques to access cytC in its natural mitochondria environment.

Despite numerous studies on isolated cytC there is a lack of understanding of conformational and functional changes of cytC and its tuning mechanisms to comply with electron flux in ETC in living mitochondria. Raman spectroscopy (RS) was used to study reduced *c*- and *b*-type cytochromes in isolated cytochromes and ETC complexes, cells, tissues, and whole organs^21,22,23,24,25,26^. However, RS cannot provide information about certain heme conformations and cannot detect oxidized cytochromes due to their weak Raman scattering. Surface-enhanced Raman spectroscopy (SERS) is known to be a highly effective tool to enhance Raman scattering of molecules^28,29,30,31,32,33^. SERS allows for the study of molecules at very low (below nM) concentrations and is widely used in biochemistry, pharmacology, and biomedical sciences^34,35^. Previously, we developed a sensitive SERS-based approach for the selective enhancement of Raman scattering from cytC heme in intact mitochondria using silver nanostructured surfaces^27^. Shin, Lee and colleagues applied triphenylphosphine-modified gold nanoparticles to enhance Raman scattering from reduced cytC in mitochondria of living cells and to monitor cytC oxidation under apoptosis and application of TNF-α^36,37^.

Here for the first time we introduce a SERS-based approach as a highly sensitive and selective tool to study the conformational changes of heme in oxidized cytC in intact functioning mitochondria and to estimate the rate of the electron transport in the respiratory chain and the overall mitochondria redox status. Our study identifies that cytC heme switches between planar and ruffled conformations in response to changes in mitochondria membrane potential and electrochemical H^+^-gradient to regulate the efficiency of the mitochondrial ETC, thus, adjusting the mitochondrial respiration to the cell’s activity.

## RESULTS AND DISCUSSION

### Assignment of SERS spectrum

For SERS measurements we used plasmonic Ag nanostructured surfaces (AgNSS) of a complex morphology allowing for strong field enhancement (Supplementary Fig. 1–4). These structures have no inhibitor effect on mitochondria respiration and ETC function (Supplementary Fig. 5 and 6).

To clarify the origin of the peaks in SERS spectra of mitochondria we compared four preparations: isolated mitochondria (Fig. 1a,b), submitochondrial particles (SMP – vesicles of inverted inner mitochondrial membrane with all ETC complexes, ATP-synthase, and cytC encased inside) (Fig. 1c,d), liposomes made of phosphatidylcholine and cardiolipin with the membrane-bound cytC (Fig. 1e,f), and purified cytC. In the mitochondria, cytC molecules are located closer to AgNSS than other ETC cytochromes (Fig. 1b). On the contrary, in the SMPs, the “matrix side” of the inner mitochondrial membrane attaches to AgNSS and, thus, *b*-type and *c*_*1*_ cytochromes are located closer to AgNSSs than cytC molecules (Fig. 1d). In the liposomes, cytC molecules were encapsulated inside liposomes and bound to the inner membrane surface (Fig. 1f). Each preparation was placed on AgNSS and illuminated with the laser light of wavelength 514 or 532 nm – pre-resonance and resonance conditions for *c*- and *b*-type hemes in cytochromes of ETC (Supplementary Fig. 7). Under these excitation conditions cytC, cytochromes *c*_*1*_ and *b*_*low*_, *b*_*high*_ in complex III and cytochrome *b* in complex II give resonance Raman scattering while *a*-type cytochromes in complex IV do not^2,22,24^. An intensive SERS spectrum of mitochondria was recorded (Fig. 1h, spectrum 1) under experimental protocol that ensures electron donation to ETC through complexes I (by malate and pyruvate) and II (by succinate), and to secure ATP synthesis (by ADP).

**Figure 1.**
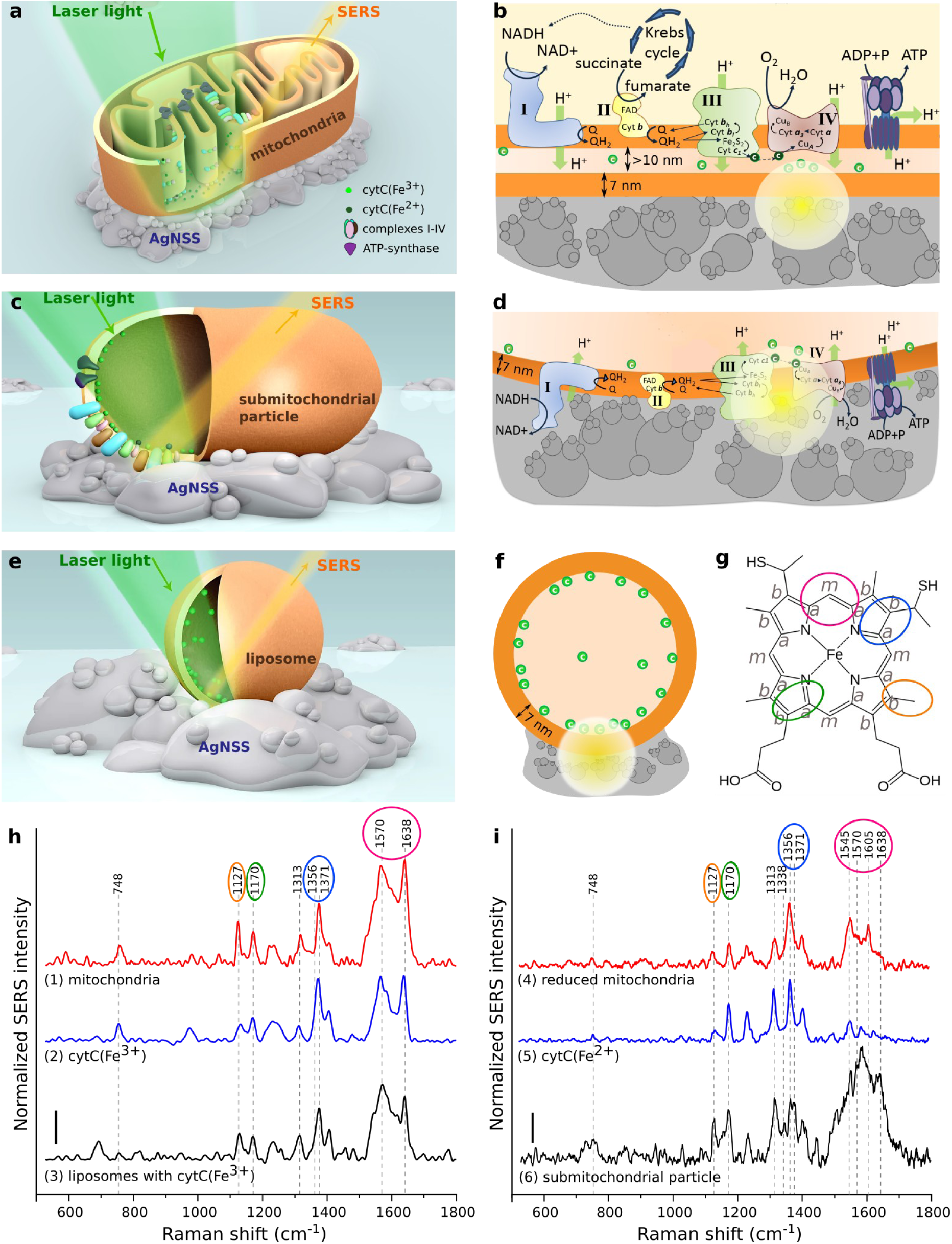
Surface-enhanced Raman scattering from cytochrome *c* molecules in mitochondria, inverted mitochondria, and liposomes. **a, b** Schematic drawing of mitochondria and its submembrane region with ETC complexes and cytochrome *c* placed on Ag nanostructured surface (AgNSS) illuminated by laser light. SERS originates from oxidized cytC molecules. Oxidized cytC molecules are marked as light green circles; reduced cytC molecules are shown with dark green circles next to complexes III or IV. **c, d** Schematic drawing of a submitochondrial particle (SMP) with ETC complexes and cytС. SERS originates from reduced and oxidized b- and c-types cytochromes. **e, f** Schematic drawing of a liposome containing oxidized cytC bound to the inner membrane surface. **g** Structural formula of a heme *c* molecule. Heme atoms and bonds of interest assessed in SERS spectra are marked in red, blue, orange, and green. **h** SERS spectra of mitochondria, purified oxidized cytC (cytC(Fe^3+^)) and cardiolipin-containing liposomes with internal membrane-bound oxidized cytC (spectra 1–3, respectively). **i** SERS spectra of mitochondria reduced with sodium dithionite, purified reduced cytC (cytC(Fe^2+^)), and submitochondrial particle (spectra 4–6, respectively). SERS spectra 1– 3 are normalized by the intensity of the peak at 1371 cm^-1^; SERS spectra 4–6 are normalized by the intensity of the peak at 1356 cm^-1^. Vertical bars in **h** and **i** correspond to 1 r.u, x-axes correspond to Raman shift, cm-1; y-axes correspond to the normalized SERS intensity.

The peak at 1313 cm^-1^ is the characteristic feature of the *с*-type heme (Fig. 1g). This peak is present only in Raman and SERS spectra of *c*-type cytochromes, but not of *b*-type cytochromes^21^. SERS spectrum of mitochondria also contains peaks at 1638 and 1570 cm^-1^, 1371 and 1172 cm^-1^, 1127 cm^-1^ and 748 cm^-1 18,21,22,23,24,27,38^. The detailed peak assignment is given in Supplementary Table 1. SERS peaks at 1371 and 1638 cm^-1^ are the signature of oxidized, but not reduced, cytochromes (Fig. 1h). We observed the characteristic peaks of reduced cytochromes at 1356, 1545, and 1605 cm^-1^ only after mitochondria treatment with sodium dithionite that reduces all electron carriers in ETC (Fig. 1i, spectrum 4). The SERS spectrum of mitochondria after sodium dithionite treatment is similar to the SERS spectrum of reduced cytC (Fig. 1i, spectra 4 and 5). SERS spectra of mitochondria do not contain the characteristic peak (1338 cm^-1^) of *b*-type cytochromes of complexes II and III. We assume that the absence of peaks assigned to b-type cytochromes and reduced cytochrome *c* is due to the fact that they are located further away from AgNSS than cytC is.

Indeed, we observed characteristic spectral features of b-type cytochromes in SMP where *b*-type cytochromes are located closer to AgNSS than cytC molecules. SERS spectra of SMP have characteristic peaks at 1313 cm^-1^ (*c*-type heme) and 1338 cm^-1^ (*b*-type heme) (Figs. 1i spectrum 6) and a peak at 1356 cm^-1^ corresponding to reduced hemes (Fig. 1i, spectra 4-6). This means that the enhancement of Raman scattering occurs from both redox forms and both *c*- and *b*-type cytochromes since they are located close to AgNSS. These experimental data confirm that SERS spectra of functional mitochondria correspond to oxidized cytC only: oxidized cytC is located closer to AgNSS than reduced cytC and oxidized/reduced cytB and cytochrome *c1* are. We suggest that oxidized cytC performs 3D diffusion in the mitochondrial IMS in contrast to reduced cytC that moves along the IMM and complexes III, IV in the supercomplex of respirasome. Such confined movement of the reduced cytC along a particular route was discussed^39^ but to the best of our knowledge, has never been observed experimentally in intact functioning mitochondria in solution.

To estimate changes in the conformation of cytC heme we used the peak at 1638 cm^-1^ – a signature of the planar heme conformation corresponding to the in-plane vibrations of methine bridges (Fig. 1g)^17,38^. Intensity of this peak increases with increasing probability of planar heme conformation and decreases with the increasing probability of ruffled heme conformation^17,38^. The peak at 1371 cm^-1^ corresponds to symmetric pyrrole half-rings ν_4_ vibrations in oxidized cytC. The intensity of this peak remains constant under changes in heme conformation and, therefore, can be used for normalization. The ratio I_1638_/I_1371_ can serve as a qualitative estimation of the probability of the planar heme conformation. We also used the ratios I_1170_/I_1371_ and I_1170_/I_1127_ to evaluate the probability of the asymmetric *vs*. symmetric pyrrole half-ring vibrations and the probability of vibrations of heme CH_3_-radicals. Previously we demonstrated that these ratios are sensitive to the conformational mobility of protein in the heme surrounding that can affect heme conformation^18^. Under applied stimuli we monitored the 1300-1400 cm^-1^ region: the showing of the peak at 1356 cm^-1^ would indicate the appearance of reduced cytochromes in the region of SERS signal registration and the showing of the peaks at 1300 and 1338 cm^-1^ would indicate appearance of *b*-type cytochromes in the region of SERS registration. Importantly, under all studied conditions we never observed the mentioned peaks demonstrating that SERS spectra of mitochondria originate from the heme of oxidized cytC and that any change in SERS peak intensity ratios corresponds to the change in the heme conformation of the oxidized cytC.

### Regulation of the cytC heme conformation in functioning mitochondria

CytC is the only one electron carrier diffusing in the intermembrane space of mitochondria, therefore it is subjected to the rapidly changed properties in its local environment. Due to ETC function cytC constantly experiences changes in the local electromagnetic field and H^+^ gradient. To find out whether the potential of inner mitochondrial membrane or H^+^ concentrational gradient influence the conformation of cytC heme, we performed series of experiments affecting ΔΨ by introducing changes in the H^+^ or K^+^ gradients across IMM.

### Proton electrochemical gradient affects conformation of cytC heme

ΔΨ is strongly influenced by the changes in the proton (H+) electrochemical gradient across the inner mitochondrial membrane created during the electron transport in ETC (Fig. 2a). To dissipate the H^+^-gradient we applied protonophore CCCP (Carbonyl cyanide *m*-chlorophenyl hydrazone) – a highly selective H^+^-ionophore that incorporates into the IMS and transfers protons from the intermembrane space to the mitochondrial matrix. This diminishes proton electrochemical gradient on IMM, increases pH of the intermembrane space, and decreases ΔΨ; the respiratory chain begins to work more intensively, transferring electrons faster in order to restore the proton gradient and inner mitochondrial membrane potential (Fig. 2a)^40^. This is manifested in SERS spectra of mitochondria through an increased intensity in the region of 1500–1700 cm^-1^ (Fig. 2b, spectrum 3) compared to the original SERS spectrum of mitochondria after substrate application (Fig. 2b, spectrum 2). An increase in the ratio I_1638_/I_1371_ after CCCP application (Fig. 2c) corresponds to the increased probability of methine bridges vibrations (C_a_C_m_, C_a_C_m_H bonds, Fig. 1g) in comparison with symmetric vibrations of pyrrole half-rings in the oxidized cytC heme. This is associated with an increased in-plane mobility of the cytC heme and a higher probability of the planar conformation of heme that favors a higher rate of electron transfer. We did not observe significant changes in the ratios I_1170_/I_1371_ and I_1170_/I_1127_. This indicates the absence of changes in vibrations of CH_3_-radicals and asymmetric vibrations of pyrrole half-rings, and hence in the conformational mobility of the proteins surrounding the heme molecule (Fig. 2d,e). CCCP did not cause changes in the ratio I_1638_/I_1371_ when applied after rotenone that prevents the accumulation of protons in the intermembrane space due to the selective inhibition of complex I (Fig. 2f): CCCP effect on the cytC heme conformation is observed only when the proton gradient across IMM is established. On the contrary, a decline in the ratio I_1638_/I_1371_ was observed after the application of oligomycin that inhibits ATP-synthase and leads to the accumulation of protons in the intermembrane space of mitochondria, to the hyperpolarization of IMM, and to the slow-down of ETC (Fig. 2g). The decreased ratio I_1638_/I_1371_ corresponds to the decreased in-plane mobility, a lower probability of the planar heme conformation, and a higher probability of the deformed, ruffled heme conformation. The subsequent application of CCCP that transfers protons from the IMS to the mitochondrial matrix and restores ETC activity led to an increase in the ratio I_1638_/I_1371_ in oligomycin-treated mitochondria (Fig. 2g). Moreover, we detected an increase in the ratio I_1638_/I_1371_ in SERS spectra of mitochondria with the increase in pH of the physiological buffer used for mitochondria (Fig. 2h). This result means that cytC heme is sensitive to electrochemical H^+^-gradient on IMM and to ΔΨ: the probability of planar heme conformation increases, when electrochemical H^+^-gradient becomes less negative and when ΔΨ decreases. (Fig. 2h). Neither rotenone, nor oligomycin, nor extramitochondrial pH affected other peak ratios in mitochondria SERS spectra (Supplementary Fig. 8). The SERS spectra of isolated purified cytC were not influenced by pH in the studied range, that can be explained by the absence of direct effect of protons on purified cytC in the solution (Supplementary Fig. 9). These results confirm that the disappearance or restoration of electrochemical proton gradient in mitochondria affects cytC heme conformation not via direct interaction of protons with cytC but due to complex processes initiated by ΔΨ changes under H^+^ movements across mitochondrial membranes. It should be noted, that in the parallel with the changes in the I_1638_/I_1371_ ratio under all studied stimuli we observed similar changes in the ratio of sum intensities in the spectral region 1530–1700 cm^-1^ (assigned to methine bridges vibrations of different symmetry types) and the spectral region 1300– 1400 cm^-1 (^assigned to various vibrations of groups in pyrrole rings) that additionally demonstrated the increased probability of the planar cytC heme conformation (Supplementary Figure 10).

**Figure 2.**
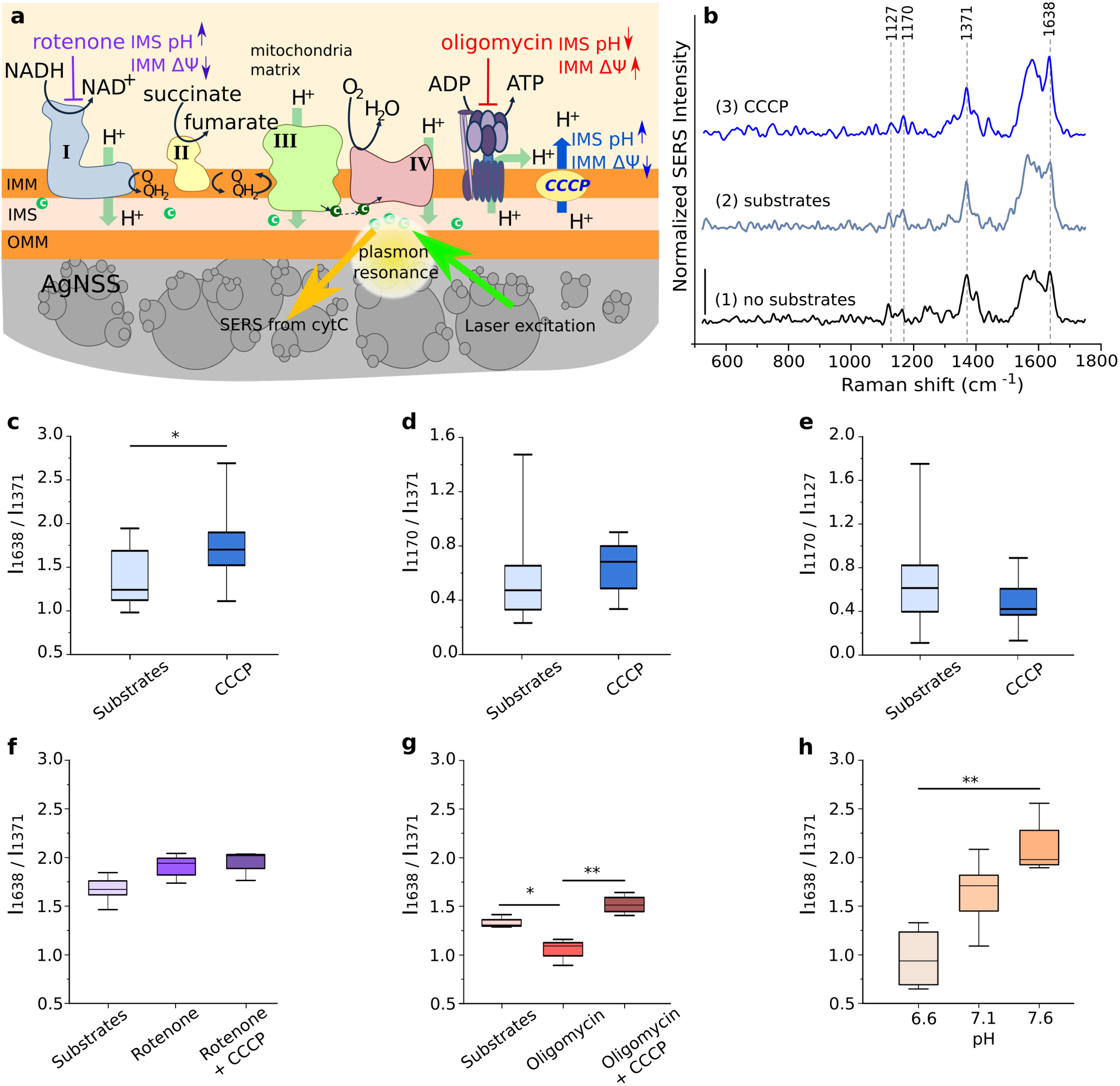
Proton gradient affects the conformation of cytC heme. **a** Schematic drawing of mitochondria on Ag nanostructured surface (AgNSS): outer and inner membranes (OMM and IMM), intermembrane space (IMS), ETC complexes, oxidized and reduced cytochrome *c* molecules (light and dark green circles in IMS), and sites of rotenone, oligomycin and CCCP action. **b** SERS spectra of mitochondria before and after application of ETC substrates (malate, pyruvate, succinate, and ADP), and after CCCP treatment (0.5 μM) (spectra 1-3, respectively). SERS spectra are normalized by the intensity of the peak at 1371 cm^-1^. Vertical scale-bar corresponds to 1 r.u. **c**–**e** The ratios I_1638_/I_1371_ (the probability of the planar heme conformation), I_1170_/I_1371_ (the probability of the asymmetric pyrrole half-ring vibrations), and I_1170_/I_1127_ (probability of vibrations of heme CH_3_-radicals) before and after CCCP treatment (n=9). **f** The ratio I_1638_/I_1371_ before and after application of rotenone (50 nM), and after the following application of CCCP (0.5 μM) (n=4). **g** The ratio I_1638_/I_1371_ before and after administration of oligomycin (10 μM), and after the following application of CCCP (0.5 μM)) (n=4). **h** Dependence of the ratio I_1638_/I_1371_ on pH in mitochondrial buffer (n=4). Data in **c-h** are represented as boxplots. All boxplots indicate median (center line), 25th and 75th percentiles (bounds of box), and minimum and maximum (whiskers). *p<0.05, **p<0.01 (Nonparametric Kruskal-Wallis test with Dunn’s multiple comparison post-test correction).

### Potassium gradient affects conformation of cytC heme

To check whether observed changes in SERS spectra of mitochondrial cytC could be caused by changes in the potential of inner mitochondrial membrane without H^+^ transport across IMM we studied the effect of K^+^-ionophore valinomycin (Fig. 3). Valinomycin incorporates into the IMM and transfers K^+^ ions from the IMS into the mitochondrial matrix down the electrochemical K^+^-gradient resulting in a decrease of ΔΨ (Fig. 3g)^41^. Valinomycin, similar to CCCP, causes an increase in the electron transport rate in ETC in order to restore ΔΨ. We observed an increase in the ratio I_1638_/I_1371_ under the application of valinomycin to the mitochondria suspension with the pyruvate, malate, succinate, and ADP (Fig. 3a, left). Valinomycin treatment of mitochondria with substrates also caused the increase in the overall intensity in the spectral region 1530-1700 cm^-1^ comparing to the spectral region 1300-1400 cm^-1^ (Supplementary Figure 10f) that additionally indicated the increased probability of the planar conformation of cytC heme. Valinomycin is known to cause the most significant change in the mitochondrial ΔΨ value compared to other ionophores or ETC inhibitors, thus affecting mitochondria SERS spectra to the largest degree^41^. Indeed, principal component analysis discriminated between SERS spectra of mitochondria without substrates and with substrates plus valinomycin application (Supplementary Figure 11). As well as other treatments, valinomycin, however, did not affect the ratios I_1170_/I_1371_ and I_1170_/I_1127_, indicating the absence of ΔΨ influence on the vibrations of CH_3_-radicals and asymmetric vibrations of pyrrole half-rings and, thus, on the conformational mobility of the protein surrounding of the heme molecule (Figs. 3 b,c, left).

**Figure 3.**
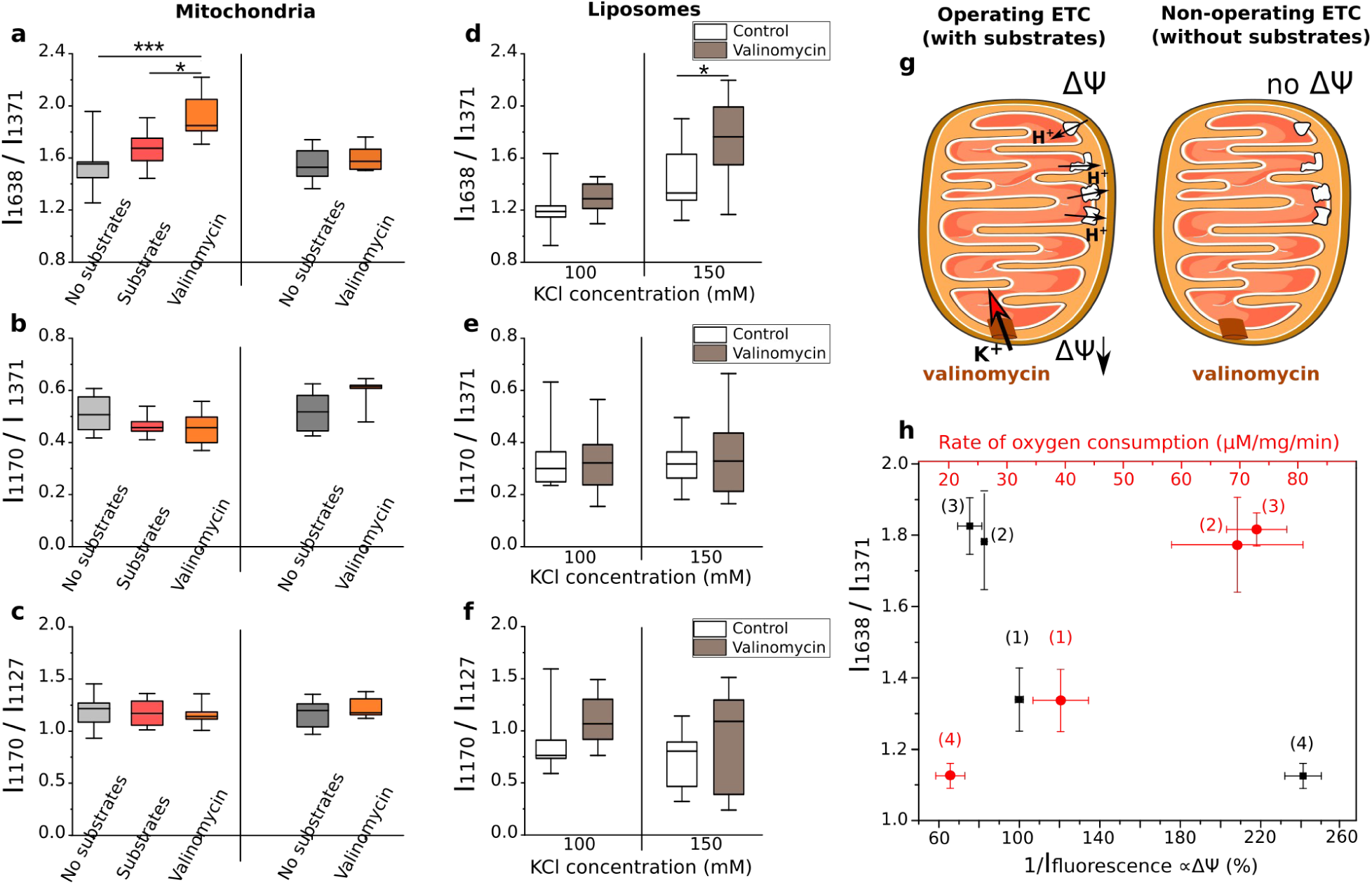
Membrane potential affects the conformation of cytC heme in mitochondria and cardiolipin-containing liposomes. **a–c** The ratios of peak intensities I_1638_/I_1371_ (the probability of the planar heme conformation), I_1170_/I_1371_ (the probability of the asymmetric pyrrole half-ring vibrations), and I_1170_/I_1127_ (the probability of vibrations of heme CH_3_-radicals) calculated from SERS spectra of mitochondria **(left panel)** with active ETC: without substrates (grey boxes), with substrates – malate, pyruvate, succinate and ADP (red boxes), and after following valinomycin application (orange boxes); **(right panel)** with non-active ETC: without substrates (dark grey boxes) and after valinomycin (0.1 μM) application (orange boxes). **d–f** The ratios I_1638_/I_1371_, I_1170_/I_1371_, and I_1170_/I_1127_ calculated from SERS spectra of liposomes before (white boxes) and after valinomycin (1 μM) application (brown boxes) for 100 and 150 mM KCl concentrations in the outside solution. Data in **a–f** are represented as boxplots. All boxplots indicate median (center line), 25th and 75th percentiles (bounds of box), and minimum and maximum (whiskers). *p<0.05, ***p<0.001 (Nonparametric Kruskal-Wallis test with post Dunns multiple comparison). **g** Schematic drawing of mitochondria with and without Krebs cycle substrates and ADP under application of valinomycin. Valinomycin provides inward K^+^ transport in mitochondria with operating ETC. **h** Ratio I_1638_/I_1371_ *vs*. the rate of oxygen consumption by mitochondria (red circles, red X-axis) and *vs*. the inner mitochondrial membrane potential ΔΨ estimated qualitatively as the reverse values of fluorescence intensity of MitoTrackerCMXRos (black squares, black X-axis). The measurements were done under the following conditions: with ETC substrates (state 1), with the sequential addition of either CCCP (state 2), or valinomycin (state 3), or oligomycin (state 4). Data are represented as mean values and errors of the mean.

Valinomycin effect on heme conformation depends on the presence of the Krebs cycle substrates and ADP that are essential for the maintenance of the electron transport in ETC and for the formation of ΔΨ on the inner mitochondrial membrane. Valinomycin transfers K^+^ ions across the IMM into the mitochondrial matrix only when ΔΨ exists (Fig. 3g). Therefore, we did not observe the effect of valinomycin on the ratio I_1638_/I_1371_ in mitochondria with non-operating ETC (without ETC substrates and ADP in the mitochondrial buffer) (Fig. 3a, right).

To test our hypothesis that CytC heme conformation depends on ΔΨ, we performed experiments with liposomes made of phosphatidylcholine and cardiolipin and containing cytC inside bound to cardiolipin (Fig. 1e,f). Before SERS measurements liposome samples were diluted with KCl solution to the final K^+^ concentrations of 100 or 150 mM. We found that SERS spectra of cytC bound to the inner surface of the liposome membrane do not depend on K^+^ concentration in the outside buffer (Fig. 3d–e, white boxes). But we observed a significant increase in the ratio I_1638_/I_1371_ under valinomycin application when the concentration of KCl in the outside buffer was 150 mM (Fig. 3d). This indicates an increased probability of the planar heme conformation. The other SERS peaks did not change (Fig. 3e–f). We did not observe any effect of K^+^ ions on SERS spectra of cytC bound only to the outer surface of the membrane of cardiolipin-containing liposomes (Supplementary Fig. 12). These results reveal that cytC heme conformation is sensitive to processes in the confined close-to-membrane spaces evoked by K^+^ transport through the membrane: changes in the local electromagnetic field, redistribution of surface charges and submembrane ions. We suggest that in mitochondria with functional ETC – where the intermembrane space is only 2-4-folds bigger than the cytC diameter and where conditions are similar to those described in the “crowding molecules” theory^42^ – the effect of ions, membrane surface charge, and local electro-magnetic field on cytC heme conformation can be greater than in liposomes and in solutions with purified isolated cytC molecules.

To demonstrate the relation between conformational changes in cytC heme and mitochondria activity we measured oxygen consumption by mitochondria under normal ETC function (with malate, pyruvate, succinate, and ADP application), under ETC activation CCCP or valinomycin application) or under ETC inhibition (oligomycin application). With the increased ETC activity (the increased oxygen consumption), SERS spectra demonstrate higher values of the ratio I_1638_/I_1371_; this corresponds to the higher probability of the planar cytC heme conformation (Fig. 3h, red symbols, states 2 and 3 compared to state 1). Under the decreased ETC activity (reduced oxygen consumption) the ratio I_1638_/I_1371_ decreases, which corresponds to the higher probability of the ruffled heme conformation (Fig. 3h, red symbols, state 4 compared to state 1). We also performed qualitative estimation of the IMM potential with the fluorescent dye MitoTrackerCMXRos to show directly correlation between ΔΨ and heme conformations. Our data confirmed that IMM depolarization evoked by CCCP or valinomycin results in the increasing probability of the planar conformation of cytC heme (Fig. 3h, black symbols, states 3 and 4 compared to state 1).

## CONCLUSION

We offered a unique SERS-based assessment of conformation changes of oxidized cytochrome *c* and their functional outcomes in intact mitochondria. To the best of our knowledge this is the first study of the link between conformation and function of cytC heme in the intact mitochondria with functioning ETC. We identified the ratio I_1638_/I_1371_ (the probability of the planar cytC heme conformation) as a reliable marker to monitor electron transport rate in ETC of functioning mitochondria. Our data revealed that under conditions of the increased ETC activity, which evokes depolarization of inner mitochondrial membrane and dissipation of electrochemical H^+^-gradient, the cytC heme has an increased in-plane mobility and higher probability of planar conformation. This favors heme *c* orientation towards heme *c1* in complex III, speeds-up ETC, and restores ΔΨ and H^+^ gradient. There are at least two possibilities how ΔΨ affects cytC heme: (i) via phosphorylation of the Tyr-residues of cytC by specific ΔΨ – dependent kinase in the IMS or/and (ii) via a direct effect of the local electromagnetic field. Several studies demonstrated the existence of ΔΨ-dependent cytC-kinase in IMS that is activated under high ΔΨ resulting in cytC phosphorylation and a decrease in cytC activity^14,15^. We suggested that phosphorylation/dephosphorylation of cytC can affect conformation of cytC heme: under high ΔΨ cytC phosphorylation results in a decrease in the probability of the planar heme conformation, whereas IMM depolarization causes a gradual dephosphorylation of cytC and the increased probability of planar conformation of cytC heme. The direct effect of ΔΨ on cytC heme can be due to the “crowding molecules” condition^.42^. Our experiments on liposomes with cardiolipin-bound cytC revealed the sensitivity of cytC heme conformation to the changes in the liposome membrane potential without any enzyme-dependent phosphorylation (Fig. 3d).

We proposed a concept that links the conformation of the cytC heme to ETC activity (Fig. 4). An increased cell activity (top panel) requires higher ATP consumption leading to a decrease in the ratio [ATP]/[ADP]. This causes activation of mitochondrial ATP synthase that utilizes H^+^ from the mitochondrial intermembrane space, which results in the dissipation of H^+^ gradient and IMM depolarization. CytC senses the IMS surrounding, which results in its heme conformational changes: the probability of the planar conformation increases. Such conformation favors its orientation towards heme in cytochrome *c1* providing electron tunneling and, thus, facilitating the electron transfer between cytochrome *c1* in complex III and cytC. This increases the rate of the overall electron transport in ETC and provides H^+^ gradient necessary for the operation of ATP-synthase and optimal synthesis of ATP to meet cellular needs for energy. Fast electron acceptance from complex III by cytC also protects mitochondria from the increased superoxide anion (O_2_^*–^) production in complex III. When the cell activity decreases (bottom panel) the regulatory cycle continues in the opposite direction.

**Figure 4.**
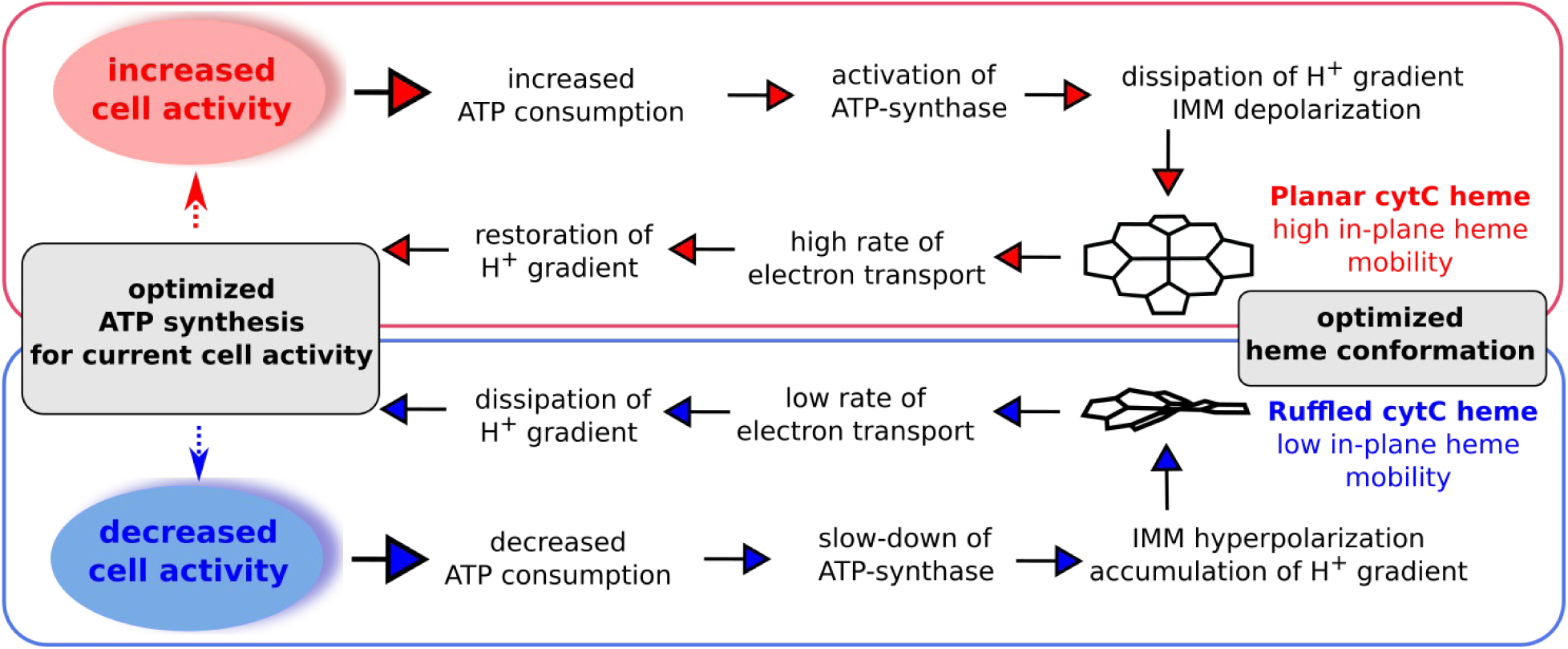
Link between cytC heme conformation and ETC activity to optimize ATP synthesis. Diagram showing the relations between cell activity, ATP consumption, inner mitochondrial membrane potential, cytC heme conformation, and ETC activity essential for the maintenance of the ATP synthesis. CytC heme can switch between planar and ruffled conformations depending on IMM potential to meet cellular needs for energy.

To conclude, the ΔΨ-regulated cytC switch between planar and ruffled heme conformations enables mitochondria to match ETC activity to the cell’s needs for ATP. The proposed SERS-based approach can provide a non-invasive, label-free tool to assess the mitochondria bioenergetics at various conditions including early-stage pathologies.

## Materials and methods

### Mitochondria isolation

Mitochondria were isolated from hearts of male Sprague-Dawley rats, body weight 275–350 g (M&M Taconic, Denmark) and male WKY rats, body weight 280–380 g (BIBCh, Pushchino, Russia)^27^. We did not observe any differences in SERS spectra of mitochondria of these strains. The animal studies were conformed with the Recommendations of the European Laboratory Animal Science Associations 2014 (FELASA) and Danish legislation governing animal experimentation, 1987, and were carried out after permission had been granted by the Animal Experiments Inspectorate, Ministry of Justice, Denmark and by Bioethics Committee of Moscow State University (protocol №82-O, 08.06.2017). Isolated mitochondria were kept in MSTP-buffer (225 mM mannitol, 75 mM sucrose, 20 mM Tris base, 2 KH_2_PO4, and 0.5 mM EDTA, pH 7.1) at 4 °C. The concentration of total protein in the obtained mitochondrial sample was 4.5–8 mg/ml. SERS experiments were performed within 2 h after mitochondria isolation.

### Liposome preparation

Liposomes with the internally bound cytC were obtained using a standard protocol: https://www.sigmaaldrich.com/technical-documents/articles/biology/liposome-preparation.html. Phosphatidylcholine (Sigma) and cardiolipin (Avanti Polar Lipids) were mixed in the ratio 4:1 by weight and diluted in chloroform in a round bottom flask to the final lipid concentration of 10 mg/ml. The solvent was removed by the rotary evaporation with a vacuum pump at 30 °C for 30 min. Lipid films were rehydrated in an equal volume of 5 mg/ml cytochrome *с* (Sigma Aldrich) in the buffer (50 mM Tris-Cl, 0.1mM EDTA, pH 7.4) at 30 °C for 30 min. The lipid-protein solution was removed from the flask after the exposure to ultrasonic bath for 5 min. The solution was then sonicated five times for 30 s with an interval of 30 s to remove lipid-protein complex aggregates. The suspension was disrupted by seven freeze-thaw cycles in liquid nitrogen followed by eleven cycles of extrusion through a filter with 100 nm pores (Avanti Mini-Extruder). Potassium chloride was added to the final concentration of 100 mM. External cytochrome *c* was removed by size exclusion chromatography with Sephadex G150 in the buffer (50 mM Tris-Cl, 0.1 mM EDTA, 100 mM KCl, pH 7.4).

Liposomes with cytC bound to the outer surface of liposome membrane were obtained by the same protocol with the only difference that lipid films were rehydrated in the equal volume of 50 mM Tris-Cl buffer with 0.1mM EDTA (pH 7.4) at 30 °C for 30 min (without cytC). After eleven cycles of extrusion through a filter with 100 nm pores cytC was added to liposomes to the final concentration of 2.2 μM and incubated at 20 °C for 30 min to insure cytC binding to cardiolipin of the outer liposome surface.

### Submitochondrial particle preparation

Inside-out submitochondrial particles (SMP or inverted mitochondria) were prepared from bovine heart, activated, and coupled by treatment with oligomycin (0.5 μg/mg of SMP protein), as described^43^. Respiratory control ratio with NADH was 6.5 indicating a functional ETC.

### AgNSS synthesis

AgNSSs were fabricated by ultrasonic silver rain procedure as described in^27,44^ with modifications. Briefly, 2.93 g of silver nitrate (Sigma, 99.999% purity) was dissolved in 100 ml of MilliQ water under constant stirring. 7 g of NaOH were diluted in 30 ml of MilliQ and poured into the AgNO_3_ solution. The obtained dark precipitate was washed three times with 100 ml of MilliQ water. 7 ml of 10% ammonia aqueous solution was added to dissolve the precipitated; an additional 25 ml volume of the 10% ammonia solution was added. The final volume of the solution was adjusted to 170 ml with MilliQ water. The temperature of the coverslips was maintained at 340 °C (magnetic stirrer with heating IKA C-MAG HS 4 digital) during the experiment and 15 min after. The solution of silver ammonia complex was sprayed onto the entire surface of coverslips (using Albedo ultrasonic nebulizer) for an hour with 5-min breaks every 3–4 min. The coverslips were stored in the dark at room temperature.

### AgNSS characterization

SEM and TEM measurements were done by NVision 40 electron microscope (Carl Zeiss) with 7 kV accelerating voltage and LEO912 AB OMEGA microscope (Carl Zeiss), respectively.

Optical characterizations were performed on a BX51 microscope (Olympus) equipped with a halogen light source, polarizers, and a fiber-coupled grating spectrometer QE65000 (Ocean Optics) with a wavelength resolution of 1.6 nm. The reflected light was collected in backscattering configuration using MPlanFL (Olympus) objectives with magnifications ×100 (NA =0.9).

In order to reveal and visualize main features of spatial distribution of hot spots, we used a scattering-type scanning near-field optical microscope (s-SNOM) (“NeaSNOM” by Neaspec GmbH) based on an atomic force microscope (AFM) that uses cantilevered Si tips covered with platinum as near-field probes^45^. The sample was scanned in a tapping mode, with the tip oscillating at the mechanical resonance frequency Ω ≈ 250 kHz with an amplitude∼ 50 nm. The tip-sample interface was illuminated under 50° relative to the sample surface by a linearly *P*-polarized tunable (1480–1625 nm) telecom laser. The experimental conditions matched numerical simulations of a strong electric near-field in a system of AgNSS^27^.

### Raman spectroscopy measurements

Confocal Raman microspectrometers InVia Raman (Renishaw, UK) with 514 nm and 532 nm lasers were used to acquire SERS spectra of mitochondria, isolated purified cytC, SMP and liposomes with cytC. Excitation and collection of SERS signal were done via objective 20x, NA 0.4 with laser power 1.5 mW per excitation spot with 0.5–1 μm diameter. Accumulation time for each SERS spectrum was 20 s.

### SERS measurements of mitochondria

Before SERS experiments the initial mitochondrial sample (30 μl) was diluted with the physiological phosphate-based solution (150 mM KCl, 5 mM Tris-HCl, 1 mM EGTA, 5 mM Na2HPO4, 1.5 mM CaCl2, pH 7.1, 4 °С); this resulted in the experimental mitochondrial sample with the total protein concentration of 0.1 mg/ml. Sodium pyruvate (2 mM), sodium succinate (5 mM), ADP (2 mM) and MgCl_2_ (3 mM) were added to ensure electron transfer in ETC and to stimulate ATP synthesis by complex V. 300 μl of the experimental mitochondrial sample were dropped on an AgNSS placed in a MatTek glass bottom dish; SERS spectra of mitochondria with substrates (pyruvate, malate, succinate, and ADP) were recorded within 30–60 s. SERS spectra of mitochondria after the application of either protonophore CCCP (0.5 μM), or K^+^-ionophore valinomycin (0.1 μM), or oligomycin (10 μM) plus CCCP (0.5 μM), or rotenone (1 μM) plus CCCP (0.5 μM) (Sigma) were recorded within 2 min.

To study the effect of H^+^ on SERS spectra of cytC in mitochondria the initial mitochondrial sample was diluted with the phosphate-based solutions with different pH (6.6, 7.1, and 7.6). We used 514 nm laser in the experiments with CCCP and pH while 532 nm laser was used in the experiments with rotenone, rotenone plus CCCP, valinomycin, oligomycin, and oligomycin plus CCCP. We demonstrated that experimental protocols and AgNSS did not affect mitochondria respiration and membrane integrity (Supplementary Fig. 5).

### SERS measurements of liposomes

To record SERS spectra of cytC bound to the inner surface of cardiolipin-containing liposomes the initial liposome samples were diluted twice with K^+^-containing phosphate buffer to obtain liposome samples with the final K^+^ concentration of 100 and 150 mM. 300 μl of the solution with liposomes were placed into a MatTek glass bottom dish with AgNSS. SERS spectra of liposomes were obtained at laser excitation wavelength of 514 nm.

### SERS and Raman measurements of purified cytC

To record SERS spectra of isolated purified oxidized cytC (Sigma) was diluted with PBS (pH 7.2) to the final concentration of 10^−6^ M. 300 μl of cytC solution were dropped into a MatTek glass bottom dish with AgNSS. SERS spectra of cytC were obtained at laser excitation wavelength of 514 nm. To record Raman spectra of isolated purified oxidized cytC at different pH values the cytC was diluted with PBS with different pH (5.0, 6.0, 7.5, and 8.0) to the final concentration of 1 mM. 300 μl of cytC solution were dropped into a MatTek glass bottom dish. Raman spectra of cytC were obtained at laser excitation wavelength of 532 nm. Excitation and collection of Raman scattering was done via objective 20x, NA 0.4 with laser power 1.5 mW per excitation spot with 0.5–1 μm diameter, spectrum accumulation time was 20 s. The study of pH effect on heme of oxidized cytC was done by means of resonance Raman spectroscopy, not SERS, to avoid possible effect of high H^+^ concentrations on plasmonic properties of AgNSSs.

### SERS measurements of submitochondrial particle

Preliminary obtained samples of SMPs were defrosted, NADH (0.1 mM) was added and 300 μl of SMP solution was placed into MatTek glass bottom dish with AgNSS. SERS spectra of SMPs were obtained at laser excitation wavelength of 514 nm.

### Spectrum analysis

SERS and Raman spectra were processed using open source software Pyraman available at https://github.com/abrazhe/pyraman. The baseline was subtracted in each spectrum. It was defined as a cubic spline interpolation of a set of knots, number and x-coordinates of which were selected manually outside any informative peaks in the spectra. Number and x-coordinates of the knots were fixed for all spectra in the study. Y-coordinates of the knots were defined separately for each spectrum as 5-point neighborhood averages of spectrum intensities around the user-specified x-position of the knot. The baseline subtraction parameters were chosen after preliminary processing of a randomly selected set of approximately 40 spectra from different preparations to ensure that all typical baseline variations were taken into account. To compare changes in SERS spectra on various stimuli we calculated the ratios of peak intensities I_1638_/I_1371_, I_1170_/I_1371_, I_1170_/I_1127_ after the baseline subtraction or the ratio of sums of the SERS intensities in the spectral regions 1300-1400 and 1530-1700 cm^-1^ (SERS Sum (SS) ratio SS_[1530-1700]_/SS_[1300-1400]_). We also applied principle component analysis to distinguish between SERS spectra of mitochondria before and after valinomycin application.

### Oxygen consumption rate measurement

Oxygen consumption by mitochondria was measured with the Clark-type electrode using oximeter Expert001.MTX (“Econics”, Russia). Mitochondria were injected into the thermostatic chamber (24 °C) with MSTP-buffer to achieve final concentration of total protein of 0.1 mg/ml. MSTP-buffer and solutions of malate, pyruvate, succinate, ADP, CCCP, valinomycin, and oligomycin were kept in open flasks at 24 °C at least one hour before the experiments. Substrates and uncoupling agents were injected into the chamber consistently when the oxygen consumption rate reached a steady state. The rate of oxygen consumption by mitochondria was measured according to Chance’s method with modifications^40^.

### Measurements of mitochondrial membrane potential

For the measurement of mitochondrial membrane potential (ΔΨ), mitochondria were stained with the potential-sensitive fluorescent dye MitoTrackerCMXRos (ThermoFisher). The MitoTrackerCMXRos dye is known to accumulate in mitochondria proportionally the value of ΔΨ with the concentration self-quenching of its fluorescence^46^. Fluorescence spectra of mitochondria were recorded by Raman microspectrometer NTEGRA SPECTRA (NT-MDT, Russia) with the laser excitation wavelength of 532 nm, grating 150 lines/mm and laser power at the exit from the objective (x5, NA 0.15) of 0.05 mW. The accumulation time of one spectrum was 10 s. The fluorescence intensity of the MitoTrackerCMXRos was determined as the intensity of the fluorescence peak at the wavelength of 610 nm.

Isolated mitochondria were diluted with MSTP-buffer containing ETC substrates and MitoTrackerCMXRos dye (100 nM). They were incubated in the dark at room temperature for 15 min. Total protein concentration in the incubated mitochondria sample was 2 mg/ml. The suspension of mitochondria was centrifuged at 7000g, 4°C for 10 min. The supernatant was removed, the sediment homogenized and resuspended with MSTP buffer containing ETC substrates without MitoTrackerCMXRos dye and centrifuged at 7000g, 4 °C for 10 min. The supernatant was removed and the pellet was resuspended in 500 µl of MSTP-buffer until a homogeneous mitochondria sample was obtained. We recorded fluorescence spectra of the mitochondrial samples with ETC substrates, with the sequential addition of CCCP (0.5 μM) or valinomycin (1 μM). To assess the effect of oligomycin on ΔΨ mitochondria were incubated in MSTP-buffer with ETC substrates, MitoTrackerCMXRos dye (100 nM) and oligomycin (10 μM) in the dark for 15 min. For the comparison, another mitochondria sample was incubated in the MSTP-buffer with ETC substrates and MitoTrackerCMXRos dye without oligomycin for 15 min. After the incubation both mitochondria samples were centrifuged two times at 7000g, 4°C for 10 min; after each centrifugation the supernatant was replaced with MSTP-buffer plus substrates plus oligomycin or with MSTP plus substrates. To estimate changes in ΔΨ the reciprocal values of the fluorescence intensity (1/I_fluor_) of mitochondria sample after the application of ionophores or inhibitors were normalized to 1/I_fluor_ value of the mitochondria sample with substrates and given in %. An increase in 1/I_fluor_ corresponds to the hyperpolarization of the inner mitochondrial membrane while the decrease corresponds to the depolarization of the inner mitochondrial membrane.

### Statistics

Data were represented as boxplots. All boxplots indicate median (center line), 25th and 75th percentiles (bounds of box), and minimum and maximum (whiskers). For the statistical analysis we applied a nonparametric Kruskal-Wallis test with Dunns multiple comparison post-test correction. The scatter plots shown on Fig.3h are represented as mean values and error of the mean.

## Data availability

The data that support the findings of this study are available from the corresponding author under reasonable request.

## ACKNOWLEDGEMENTS

This research has been supported by the Interdisciplinary Scientific and Educational School of Moscow University “Molecular Technologies of the Living Systems and Synthetic Biology”. NAB acknowledges support from Russian Foundation for Basic Research (RFBR, grant number 20-04-01011а), EIN acknowledges support from Russian Science foundation (RSF, grant number 21-74-00026), AAS acknowledges support from Russian Science Foundation (RSF, grant number 20-73-00257). AVA and SMN acknowledge support from Russian Foundation for Basic Research (RFBR, grants number 20-07-00840a and 20-07-00475a). VSV acknowledges the financial support from the Ministry of Science and Higher Education of the Russian Federation (Agreement No. 075-15-2021-606). Scanning near field microscopy studies (Y.D.I. and A.A.V.) were funded by the Russian Science Foundation (RSF, grant number 21-72-10163). GVM acknowledges support from Russian Science Foundation (RSF, grant number 19-79-30062). We are grateful to Olga Eremina for the assistance in AgNSS preparation and to Alexey Brazhe for the help with SERS and Raman spectra analysis.

## AUTHOR CONTRIBUTION

N.A.B, E.I.N., G.V.M., E.A.G., O.S. and A.B.R. conceived the experiments. N.A.B., E.I.N., S.M.N. and O.S. performed SERS experiments and analyzed SERS spectra; A.A.S. and E.A.G. designed AgNSSs; A.A.S., E.A.G. and E.I.N. fabricated AgNSSs. N.A.B., E.I.N., Z.V.B. and A.A.B. prepared mitochondria samples; E.I.N. and V.G.G prepared liposomes with cytC; V.G.G. prepared inverted mitochondria; E.I.N., Z.V.B. and A.A.B. performed respirometry experiments and their analysis. A.A.S. and E.A.G. performed the electron microscopy (SEM) analysis. S.M.N. conducted optical spectroscopy. V.S.V., D.I.Y. A.B.E. and A.V.A. performed SNOM characterization. N.A.B., V.G.G., A.V.A., V.S.V., E.A.G., G.V.M., O.S. and A.B.R. contributed tools and materials. A.B.R. and O.S. supervised the project. N.A.B., E.I.N., S.M.N. and O.S. drafted the manuscript. All authors discussed the results, edited and commented on the manuscript.

Supporting Information Available:The following files are available free of charge: SERS_cytC_membrane_ACSSensors_revised.docx. File with the supplementary information includes Supplementary Figures 1-12 and Supplementary Table 1.

