## Supplementary Figures 1-12, Supplementary Table 1 for "SERS uncovers the link between conformation of cytochrome *c* heme and mitochondrial membrane potential"

[#] shared co-supervision

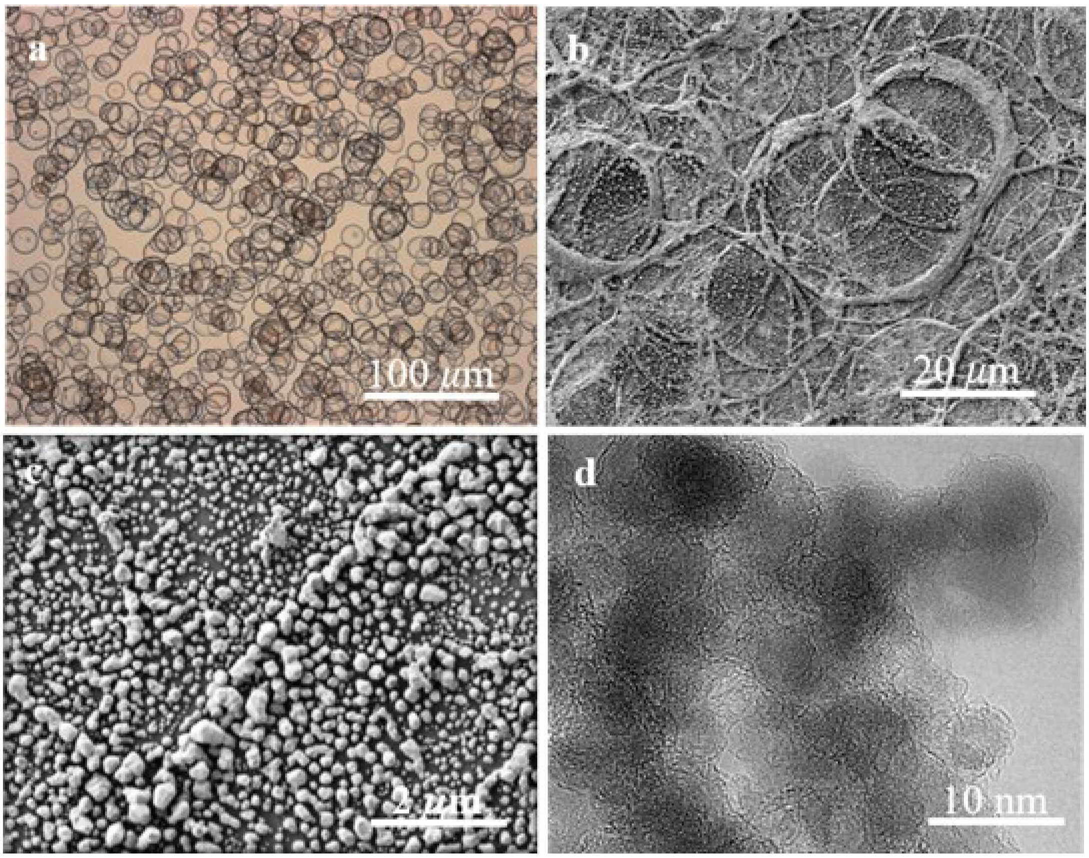


**Supplementary Figure 1.** Hierarchical structure of "coffee ring" nanostructured surfaces (AgNSS) fabricated by means of aerosol deposition. **a** Optical image of AgNSS. **b** SEM image shows structural features of silver rings. **c** Magnified SEM image contours silver clusters inside the ring. **d** TEM image reveals 2–5 nm silver nanoparticles as the smallest building blocks of the ring structure. The aerosol deposition enables the fabrication of scalable planar nanostructures with large numbers of hot spots.

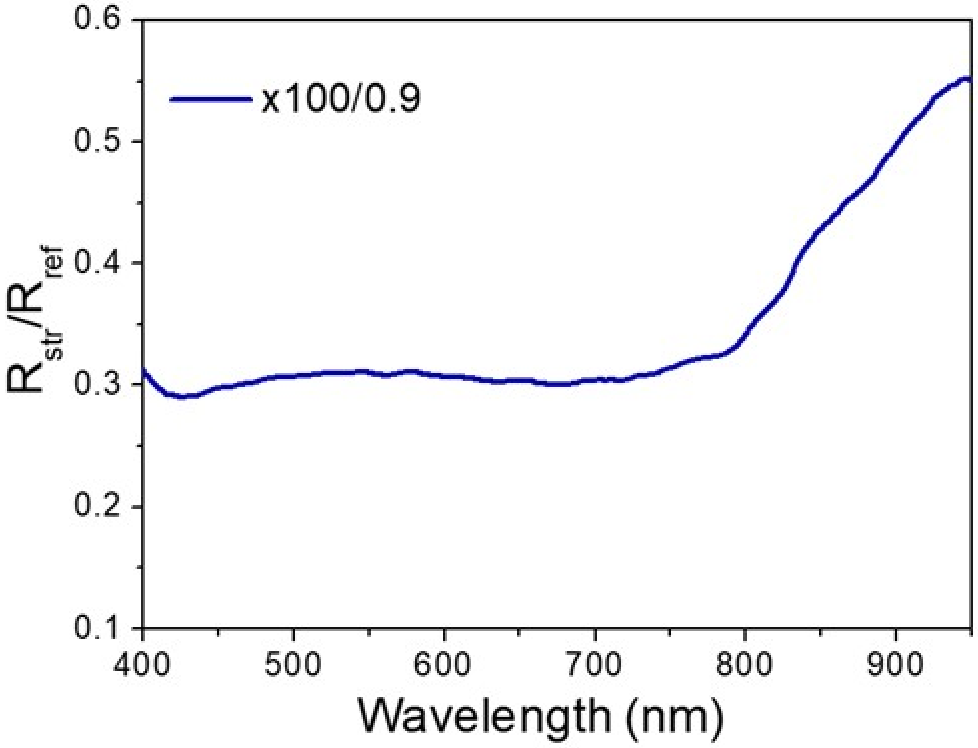


**Supplementary Figure 2.** Reflection spectrum recorded from an Ag nanostructured surface (AgNSS). The experimental data show the reflection ratio Rstr/Rref, where Rstr is the reflection measured from the AgNSS and Rref is the reference spectrum recorded from the reference broadband laser mirror, which provides an average reflection of 99% in the spectral region 350–1100 nm. AgNSSs have a very wide light absorption band due to the hierarchical nature of the nanosurfaces; this allows for their effective uses for the laser excitation in the visible and near-infrared regions.

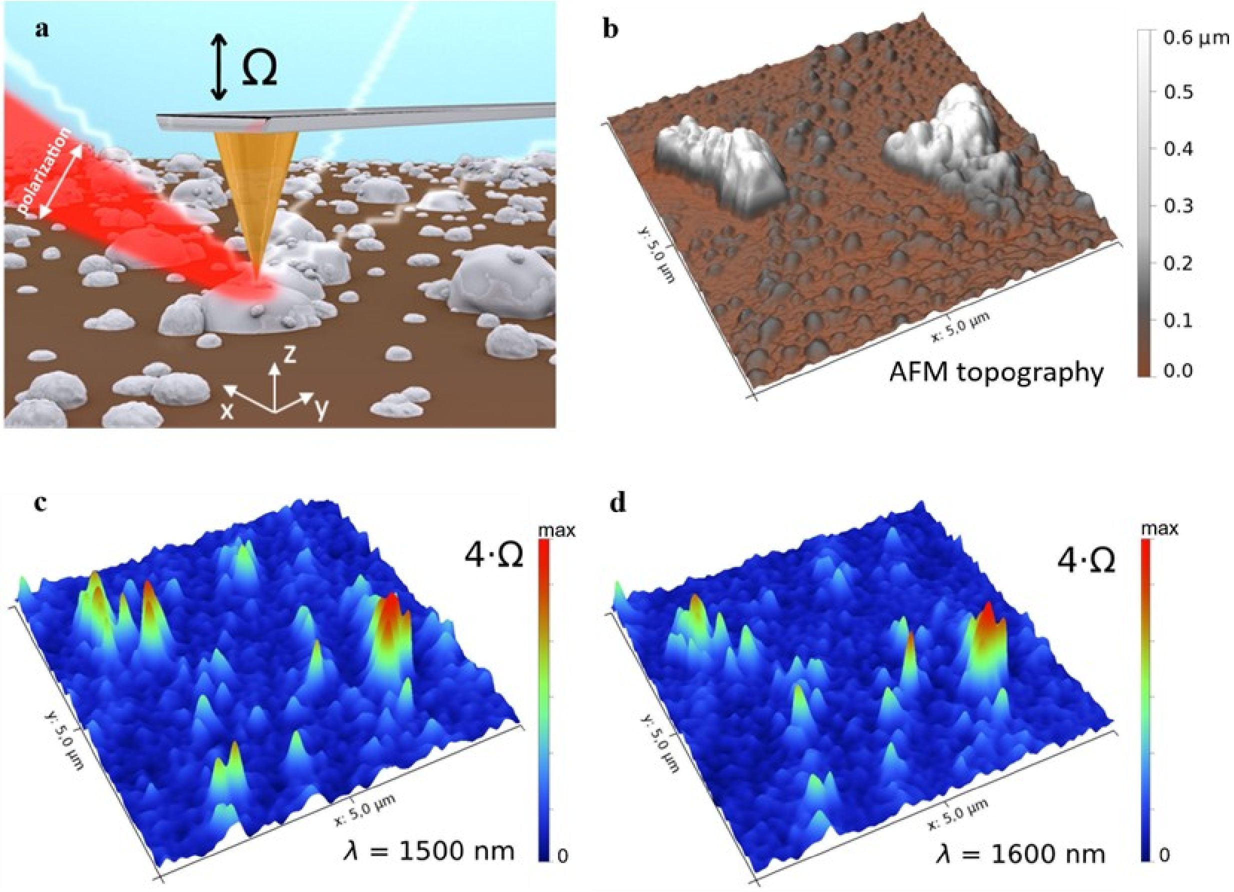


**Supplementary Figure 3**. **a** Schematic representation of particle-tip scattering in scattering-type scanning near-field optical microscopy (s-SNOM). In oblique-incidence mode, the incident polarized and focused laser beam illuminates a conglomeration of Ag nanoparticles from the top with respect to the out-of-plane axis z. The near-field scattered by the tip is collected by the detectors in the backscattered direction of propagation and acquired by a photodetector. **b** Topographical (AFM) image of Ag aggregates. **c**, **d** Typical pseudo-colour SNOM images (6 × 5 µm$^2$) of AgNSS obtained at **c** λ≈1500 nm and d λ≈1600 nm. The color scale shows optical near-field intensity. In our s-SNOM setup, optical background contributions were suppressed by demodulation of the detector signal at a high harmonic frequency of the tip oscillation nΩ (n = 4 in our case). Due to a dominant contribution of the dipole moments of nanoparticles along the z-axis, the recorded SNOM images represent mostly a distribution of the amplitude of the z-component of the electric field $E_z$. In order to enhance this selectivity, a polarizer at the detector was set correspondingly to the z-polarization of the light scattered by the tip. SNOM images reveal the existence of randomly distributed and strongly localized electromagnetic excitations (i.e., hot spots). SNOM images obtained at different illumination wavelengths (**c** and **d)** indicate that the distribution of the field enhancement weakly depends on the wavelength, and is mainly determined by the distributions of nanoparticles. This result explains why both lasers (514 and 532 nm) used in SERS measurements of mitochondria were equally effective (Supplementary Fig. 7).

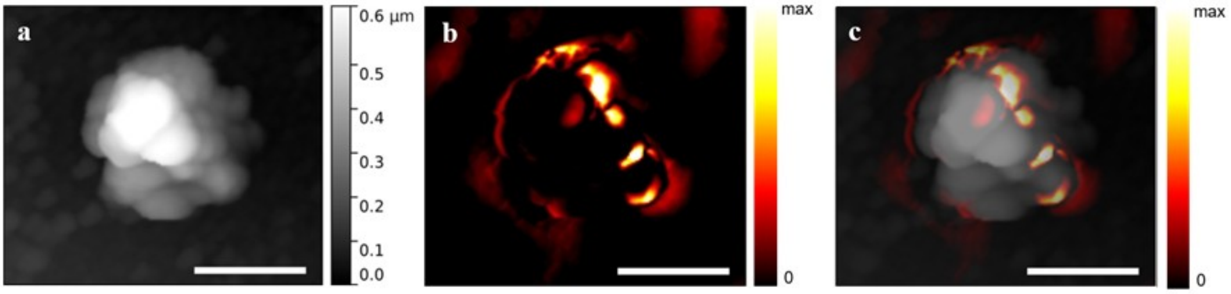


**Supplementary Figure 4. a** High resolution topographical AFM image of a typical conglomeration of silver nanoparticles on AgNSS. **b** pseudo-color SNOM image recorded at λ≈1500 nm; **c** Superimposed image of pseudo-color SNOM image (**b**) and the corresponding AFM topographical image (**a**). The localization of several hot spots is clearly seen in conglomeration of Ag nanoparticles; their appearance can be directly attributed to the near-field response of small-size nanoparticles. The strong fields (hot spots) are realized in the system providing long-range near-field distribution. This ensures that such nanostructures can be effectively used in studies of the mitochondrial respiratory chain. The scale bar is 1 $\mu$m.

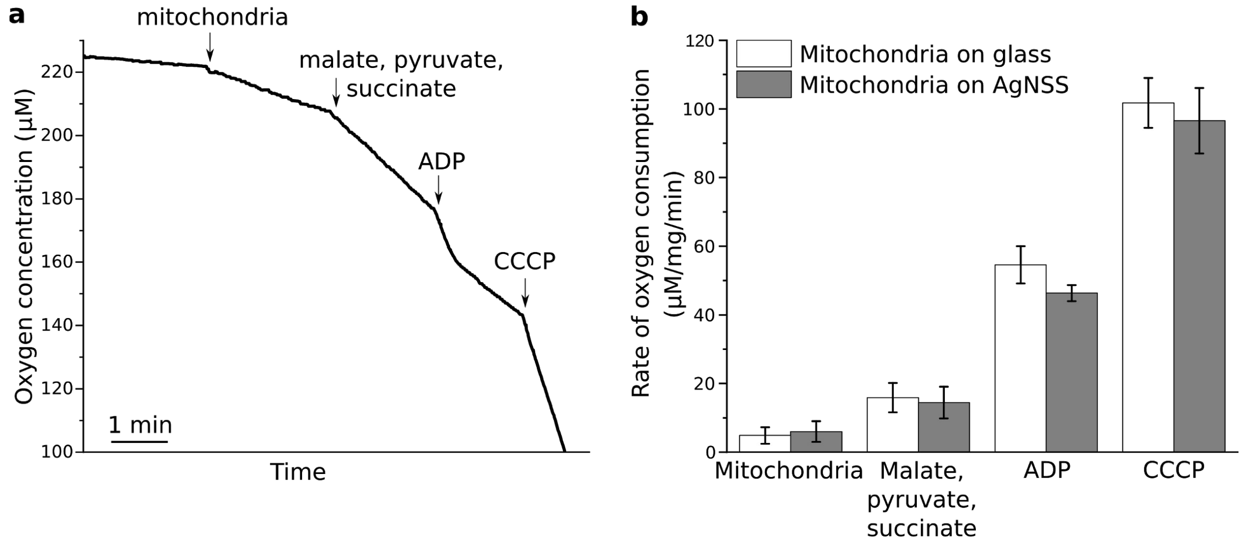


**Supplementary Figure 5. AgNSS does not affect mitochondrial respiration. a** Typical experimental curve of oxygen consumption by mitochondria under application of malate, pyruvate and succinate (to ensure electron donation to ETC complexes I and II), ADP, and protonophore CCCP. **b** The rate of oxygen consumption by mitochondria under application of malate, pyruvate and succinate, ADP, and CCCP. Mitochondrial suspensions were placed on an ordinary glass surface (white) or on AgNSS (gray) for 10 min before their injection into the experimental chamber with Clark electrode.

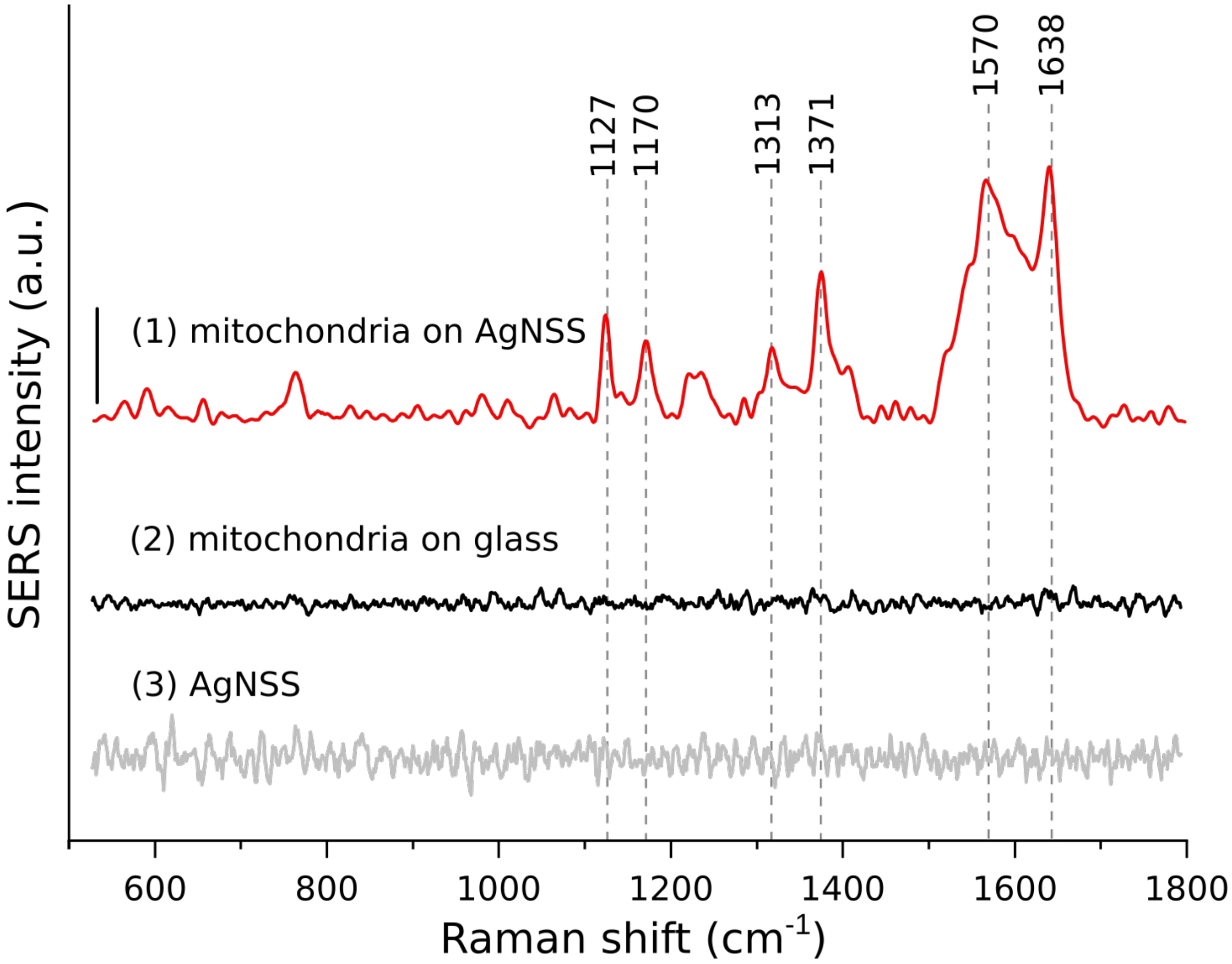


**Supplementary Figure 6.** SERS signal from mitochondria. Suspension of mitochondria placed on AgNSS gives a well-pronounced SERS spectrum (spectrum 1) in comparison to the suspension of mitochondria placed on an ordinary glass surface (spectrum 2). AgNSS in a physiological buffer without mitochondria does not give any enhancement (spectrum 3). All measurements were performed with 514 nm laser. SERS spectra are normalized by the sum of intensities of the corresponding spectra. For better representation, the spectra are shifted vertically. Vertical bar is equal to 0.002.

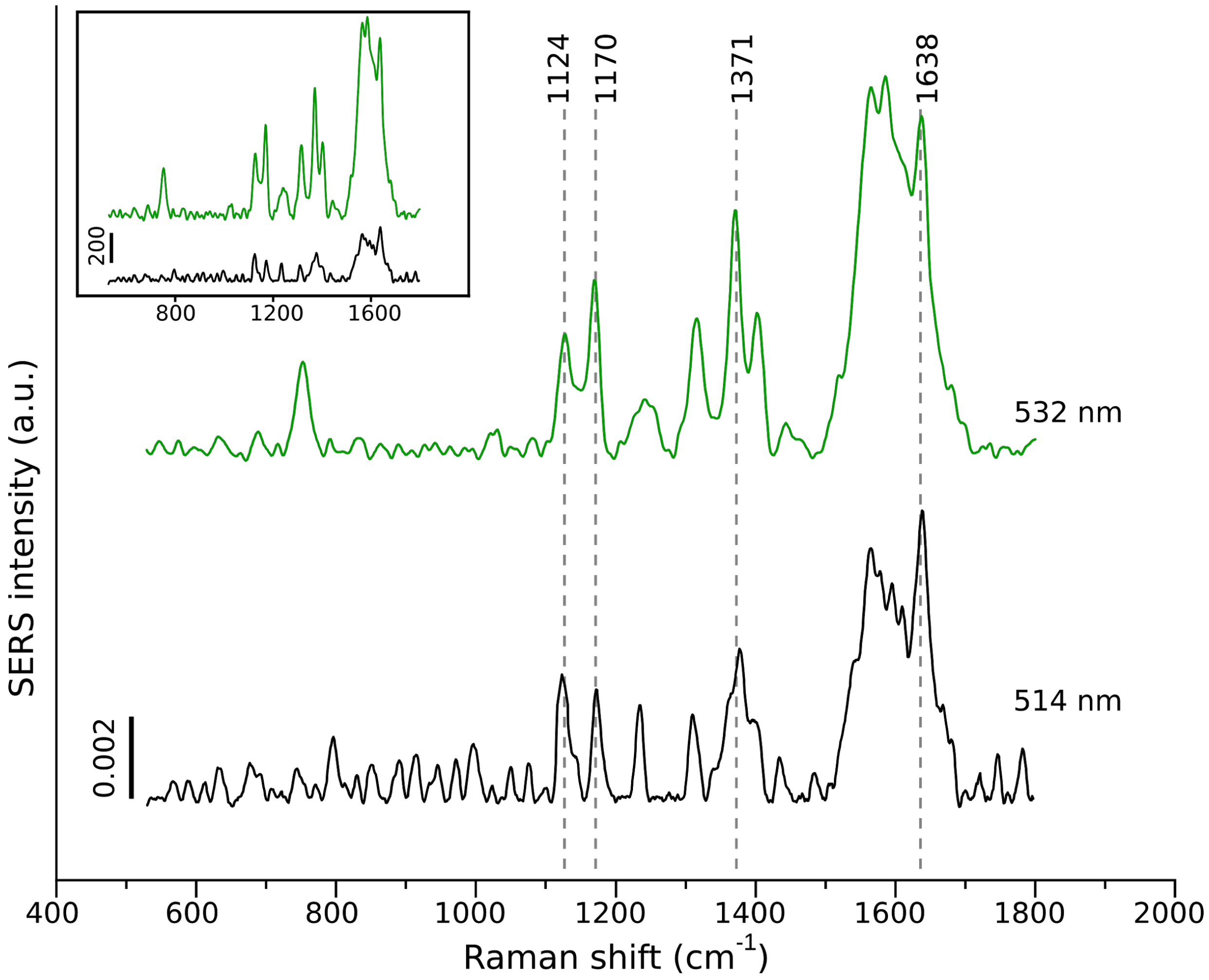


**Supplementary Figure 7.** SERS spectra of mitochondria with malate, succinate, pyruvate, and ADP obtained with 514 and 532 nm lasers. The spectra are normalized by the sum of intensities of the corresponding spectra. Inset: the same spectra without normalization.

**Supplementary Table 1.** Assignment of peaks in SERS spectra of mitochondria with ETC substrates (malate, pyruvate, succinate and ADP), SDT-treated mitochondria (fully reduced ETC), oxidized and reduced isolated cytC, liposomes with oxidized cytC, and submitochondrial particles (SMPs) with NADH.

| **Mitochondria** | | **Isolated cytC** | | **Liposomes with cytC($Fe^{3+}$)** | **SMPs +NADH** | **Vibration symmetry** | **Bond vibration** | **Sensitivity** |
|---|---|---|---|---|---|---|---|---|
| **substrates** | **SDT** | **Oxidized** | **Reduced** | | | | | |
| 1638 | 1605 | 1638 | 1608 | 1638 | 1638, 1605 | B1g, ν10 | $C_aC_m$, $C_aC_mH$, $C_aC_b$ | Planar/ruffled heme conformation, spin state of heme Fe |
| 1570 | 1545 | 1570 | 1545 | 1570 | 1570, 1545 | A2g | $C_aC_m$, $C_aC_mH$, $C_aC_b$,$C_aN$ | Spin and redox states of heme Fe |
| 1371 | 1356 | 1371 | 1356 | 1371 | 1371, 1356 | A1g, ν4 | Symmetric pyrrol half-ring | Redox state of heme Fe |
| | | | | | 1338 | | All bonds of heme *b* | The signature peak of *b*-type cytochromes |
| 1313 | 1311 | 1313 | 1311 | 1313 | 1313 | | All bonds of heme *c* | The signature peak of *c*-type cytochromes |
| 1170 | 1170 | 1170 | 1170 | 1170 | 1170 | B2g, ν30 | Asymmetric pyrrol half-ring | |
| 1127 | 1127 | 1127 | 1127 | 1127 | 1127 | B1g, ν5 | $C_b$-$CH_3$ | |
| 748 | 748 | 748 | 748 | 748 | 748 | B1g, ν15 | Heme breathing | |

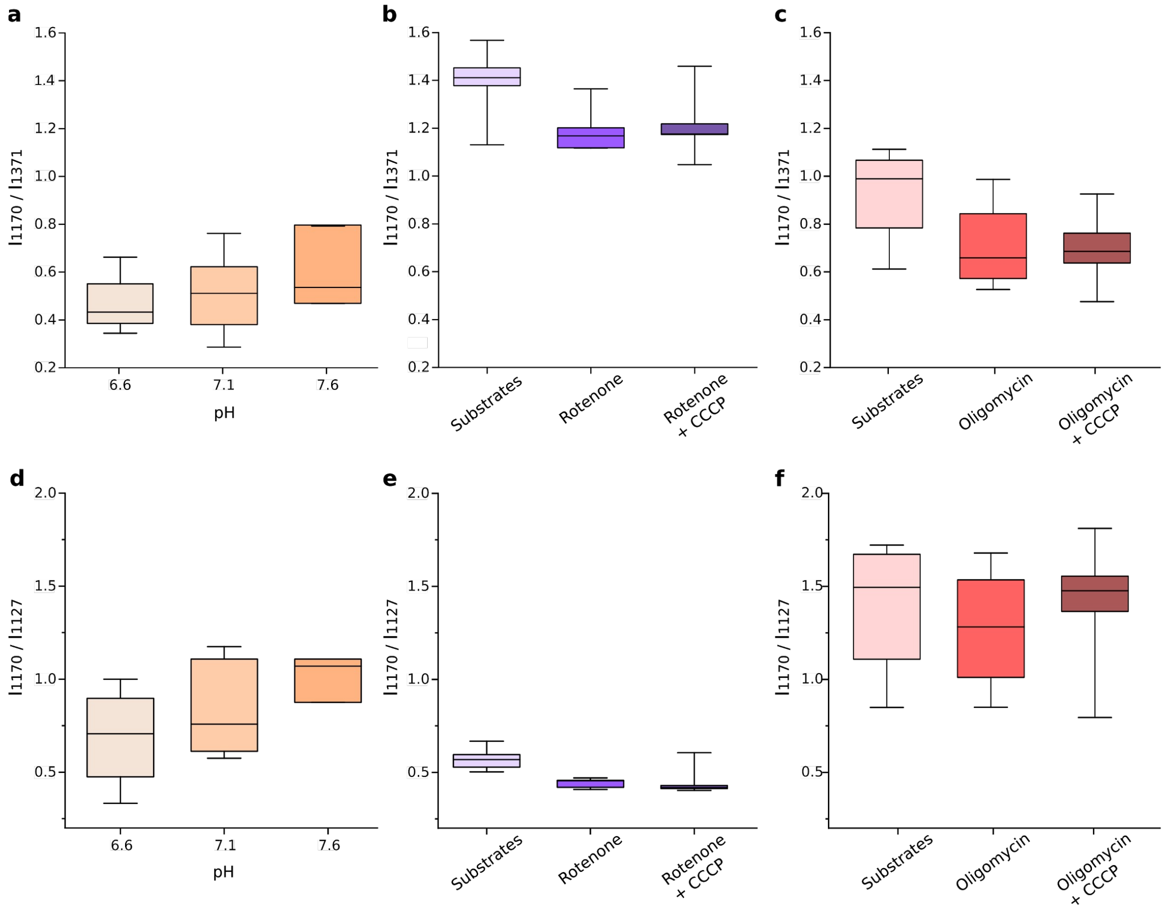


**Supplementary Figure 8.** Asymmetric vibrations of pyrrol half-rings (the ratio $I_{1170}/I_{1371}$) and vibrations of $CH_3$-radicals of cytC heme (the ratio $I_{1170}/I_{1127}$) are not affected by pH **(a, d)**, by rotenone with the sequential addition of CCCP **(b, e)**, and by oligomycin the sequential addition of CCCP **(c, f)**.

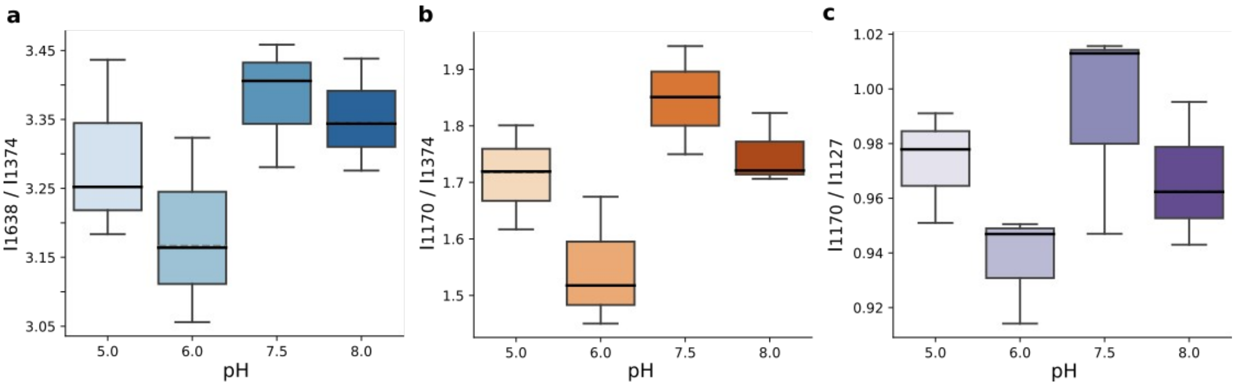


**Supplementary Figure 9.** Resonance Raman experiments on purified oxidized cytC. Heme conformation of purified oxidized cytC does not depend on pH of the buffer. The ratios $I_{1638}/I_{1374}$ (**a**), $I_{1170}/I_{1374}$ (**b**), and $I_{1170}/I_{1127}$ (**c**) are not affected by the change in pH, which indicates the absence of changes in the probability of the planar heme conformation, asymmetric vibrations of pyrrol rings, and vibrations of $CH_3$-radicals. To estimate effect of pH on the heme of purified cytC within a wide range of pH (5–8 values) we recorded resonance Raman spectra of 1 mM cytC solution placed on the ordinary glass Petri dishes (laser excitation 514 nm, obj x20, NA 0.4, 0.5 mW laser power per registration spot, 30 s of spectrum accumulation). We did not use AgNSS to avoid possible effects of acidic and basic buffers on AgNSS. The peak corresponding to the symmetric pyrrol half-ring vibrations in resonance Raman spectra is shifted to 1374 $cm^{-1}$ compared to the 1371 $cm^{-1}$ peak in SERS spectra. Such a difference is well-known for biomolecules.

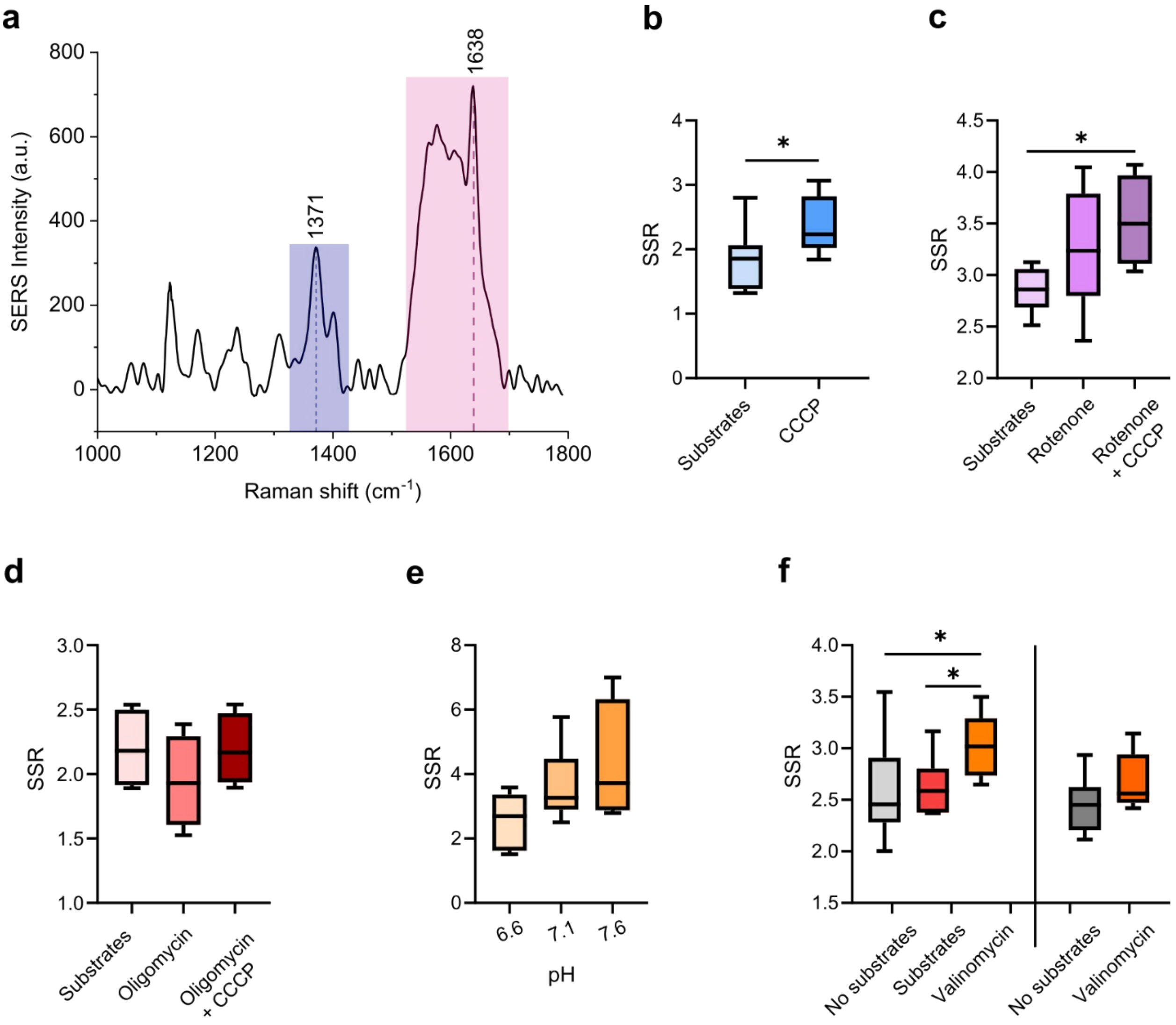


**Supplementary Figure 10. a.** SERS spectra of mitochondria with ETC substrates (malate, pyruvate, succinate, and ADP). Pink rectangle shows the spectral region characteristic to methine bridges vibrations ($C_aC_m$ bonds) of different symmetry types (1530–1700 $cm^{-1}$); blue rectangle shows the spectral region characteristic to various vibrations of $C_aC_b$, $C_aN$ groups in pyrrole rings (1300–1400 $cm^{-1}$). Figures **b-f** demonstrate boxplots for the ratio of sums of the SERS intensities in the specified spectral regions (SERS Sum Ratio (SSR) – the ratio $SS_{[1530-1700]}/SS_{[1300-1400]}$) before and after different treatments. As well as $I_{1638}/I_{1371}$ ratio, the SERS sum ratio is sensitive to the probability of the planar conformation of the cytC heme. The bigger the ratio is, the bigger the probability of the planar heme conformation is. **b**: application of $H^+$-ionophore CCCP; **c**: consequent application of rotenone and CCCP; **d**: consequent application of oligomycin and CCCP; **e**: effect of the mitochondrial buffer pH; **f**: application of $K^+$-ionophore valinomycin with and without ETC substrates. All boxplots show median (central horizontal line), 25th and 75th percentiles (bounds of box), and minimum and maximum values (whiskers). *$p<0.05$ (Nonparametric Kruskal-Wallis test with Dunn's multiple comparison post-test correction). The results are almost the same as the results obtained by calculating the ratio $I_{1638}/I_{1371}$ for the studied stimuli (Figure 2 c, f-h; Figure 3 a).

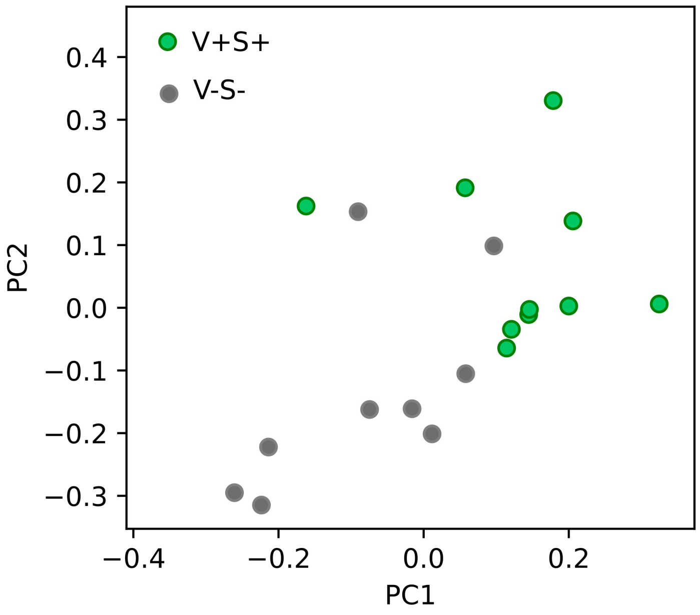


**Supplementary Figure 11.** Discrimination of the SERS spectra of mitochondria without substrates (gray dots) and with substrates after valinomycin application (green dots) using principal component analysis (PCA). Figure shows PCA score plot on the first two PC dimensions obtained with the standard PCA approach.

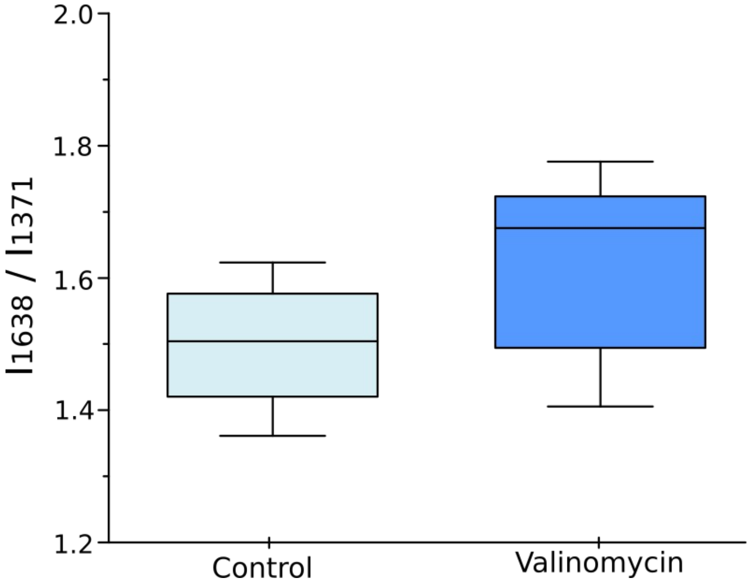


**Supplementary Figure 12.** Valinomycin does not affect the probability of the planar heme conformation (the ratio $I_{1638}/I_{1371}$) of cytC molecules bound to the outside surface of cardiolipin-containing cytochromes. Concentration of KCl in the liposome buffer  is 150 mM.
